# Virome-wide serological profiling reveals association of herpesviruses with obesity

**DOI:** 10.1101/2020.09.02.280503

**Authors:** Mohammad Rubayet Hasan, Mahbuba Rahman, Taushif Khan, Amira Saeed, Sathyavathi Sundaraju, Annaliza Flores, Philip Hawken, Arun Rawat, Naser Elkum, Khalid Hussain, Rusung Tan, Patrick Tang, Nico Marr

**Affiliations:** Department of Research, Sidra Medicine, Doha, Qatar; Department of Pathology, Sidra Medicine, Doha, Qatar; Division of Endocrinology, Department of Pediatrics, Sidra Medicine, Doha, Qatar; Weill-Cornell Medical College, Doha, Qatar; College of Health and Life Sciences, Hamad Bin Khalifa University, Doha, Qatar

**Keywords:** Obesity, seroprevalence, herpesviruses, herpes simplex virus, picornaviruses

## Abstract

The relationship between viral infection and obesity has been known for several decades but epidemiological data related to obesity is limited to only a few viral pathogens. To identify associations between viral infections and obesity, a high-throughput virome-wide serological profiling tool, VirScan, was used to measure antibody responses to a wide range of viruses. Serum specimens from 457 Qatari adults (lean=184;obese=273) and 231 Qatari children (lean=111;obese=120) were assessed by VirScan. Pediatric specimens were simultaneously tested by conventional serology for several herpesviruses to validate VirScan results. Viral association with obesity was determined by calculation of odds ratio (OR) and *p*-values from Fisher test, and by multivariate regression analysis to adjust for age and gender, with Bonferroni correction for multiple testing. Comprehensive serological profiling of Qatari adult population with VirScan revealed positive and negative associations (*p*<0.05) of antibody responses to members of Herpesviridae and Picornaviridae families, respectively, with obesity. After adjusting *p*-values for multiple comparisons, only herpes simplex virus 1 (HSV-1) and Rhinovirus A were positively (OR=3.3; 95%CI 2.15-4.99; *p*=2.787E-08) and negatively (OR=0.4; 95%CI 0.26-0.65; *p*=1.175E-03) associated with obesity. At the peptide level, higher prevalence of antibodies against several peptide epitopes of HSV-1/2 was positively (OR=2.35-3.82; *p*≤3.981E-05) associated with obesity. No such associations were seen at the species or peptide levels in the pediatric population. By multivariate regression analysis, HSV-1 was independently associated with obesity irrespective of age and gender. These findings are in agreement with limited data on the adipogenic properties of HSV-1 observed in vitro.

**Importance:** The state of Qatar has one of the highest rates of obesity and associated morbidities in the world. Although obesity is predominantly caused by the intake of high calorie diet and reduced physical activities, other factors including infections with certain viruses have been reported. Among these viruses, human adenoviruses were widely studied but epidemiological data for other viruses in relation to human obesity are limited. Here, we studied the association of obesity in Qatari adults and children with a wide range of viral pathogens using VirScan, a virome-wide serological profiling tool. Our results indicate significant association HSV-1 with obesity in the adult population only. Furthermore, we have identified a set of HSV peptides as candidate obesogenic factors for future studies.

## Introduction

The increasing prevalence of obesity has led to an increased prevalence of many chronic diseases including diabetes, heart disease, cancer and mental health conditions. A recent Global Burden of Disease (GBD) study revealed that high BMI accounts for 4 million deaths worldwide, 60% of which occurred among obese persons. Leading causes of death and disability related to high BMI include cardiovascular disease, diabetes, chronic kidney disease, cancer and musculoskeletal disorders (1-3). The rate of obesity in Qatar is one of the highest in the world. Based on the recent data published by the Qatar Biobank, 48% of Qatari men and 40% of Qatari women are obese. The high rate of obesity is also correlated with the fact that 44% of all participants have elevated cholesterol and 15.5% had previously been diagnosed with diabetes mellitus (4, 5). Although the ‘energy imbalance’ arising from increased consumption of high calorie diets and a concomitant decrease in energy expenditure due to a sedentary lifestyle is considered to be the most important cause of obesity, there are suggestions for many other contributing factors, including infectious causes (6). Evidence for infectious causes of obesity came from epidemiological links and the observation of experimental lab animals that gained body fat following infection with certain infectious agents (7).

The association between infection and obesity has been known for more than three decades and has led to the term “infectobesity” (8). To date, several viruses including adenoviruses (Adv), cytomegalovirus (CMV), herpes simplex virus 1 (HSV-1), human herpes virus 8 (HHV-8), hepatitis C virus (HCV), canine distemper virus (CDV), rous-associated virus-7 (RAV-7) and borna disease virus (BDV) have been reported to cause obesity in animals. However, epidemiological data to link infection with human cases of obesity was mostly limited to adenoviruses. Among different serotypes of adenoviruses, evidence for adipogenesis, based on results from laboratory, animal or epidemiologic investigations, exists for Adv-5, Adv-9, Adv-31, Adv-36 and Adv-37 (7, 9-12).

Despite high rates of obesity, no data on infectious causes of obesity is yet available for Qatar. While there are few reports on adenoviral infections implicated in respiratory and gastrointestinal infections, their serotypes remained unknown and none were studied in relation to obesity (13, 14). Apart from adenoviruses, laboratory and animal data suggest that other viruses may also be epidemiologically linked to human obesity. Furthermore, many other adipogenic and/or obesogenic viruses likely have not been identified yet because of the fact that seroepidemiological studies in relation to obesity were focused on specific viruses only. Studies investigating the association between adenoviruses and obesity so far were reliant on methods such as ELISA or serum neutralization assays to detect antibodies against specific viruses or on PCR methods to detect adenovirus DNA in adipose tissues (7, 15). A comprehensive seroepidemiological study may reveal the history of infection of an individual for a wide-range of pathogens and its association with the onset of obesity.

‘VirScan’ first described by Xu *et al*. (16) is a revolutionary new technique for the comprehensive serologic profiling of the human population, and can reveal the history of infections in humans. The technique is based on phage immunoprecipitation sequencing (PhIP-seq) technology that uses a bacteriophage library that displays proteome-wide peptides from a large number of human-pathogenic viruses (16). The expanded VirScan library contains approximately 115,753 56-mer peptides representing most known pathogenic, human viruses (∼400 species and strains) as well as other non-viral antigens retrieved from National Institute of Allergy and Infectious Diseases (NIAID) Immune Epitope Database (www.iedb.org) (17). To perform the serological screening, serum samples are mixed with the library allowing antibodies to bind with pathogen specific epitopes displayed on the phage surface. The bacteriophage-antibody complexes are then immunoprecipitated and the phage DNA region encoding the artificially expressed peptide antigens to which an antibody was bound are sequenced by NGS to reveal the repertoire of anti-viral antibodies in a given serum sample. With its ability to correctly detect frequently encountered anti-viral antibodies and higher sensitivity and specificity (≥95%) with reference to standard ELISA and Western blot assays, VirScan has become a powerful new technique for high-throughput serological screening (16, 18). In this study, we employed VirScan to compare the antiviral antibody repertoires in the Qatari obese and lean population with the aim to test for associations between obesity and antiviral antibody responses at the species and peptide epitope levels.

## Results

### Participant characteristics

Serum specimens from two independent cohorts comprised of mostly Qatari nationals were assessed. The adult cohort includes a total of 457 subjects selected from 800 individuals based on BMI (WHO classification criteria; lean = BMI ≤ 25; obese = BMI ≥ 30) (Table 1). These individuals represent general Qatari population who volunteered to contribute to Qatar Biobank (QBB) - a national repository of biological specimens and health information in Qatar established to facilitate medical research on prevalent health issues (5). Average age of lean and obese subjects in the adult cohort were 33.5±12.1 and 44.7±11.9 years, respectively. Within the lean and obese groups 72.3 % and 71.8 % subjects were females, respectively. Pediatric cohort includes 231 subjects who were classified as lean (BMI 5^th^ to <85^th^ percentile) or obese (BMI ≥95^th^ percentile) according to the definitions of Center for Disease Control and Prevention (CDC). Average age of lean and obese subjects in this cohort were 11.2±2.4 years and 12.1±2 years, respectively. Within the lean and obese groups 58.6% and 43.3% subjects were females, respectively.

**Table 1:**
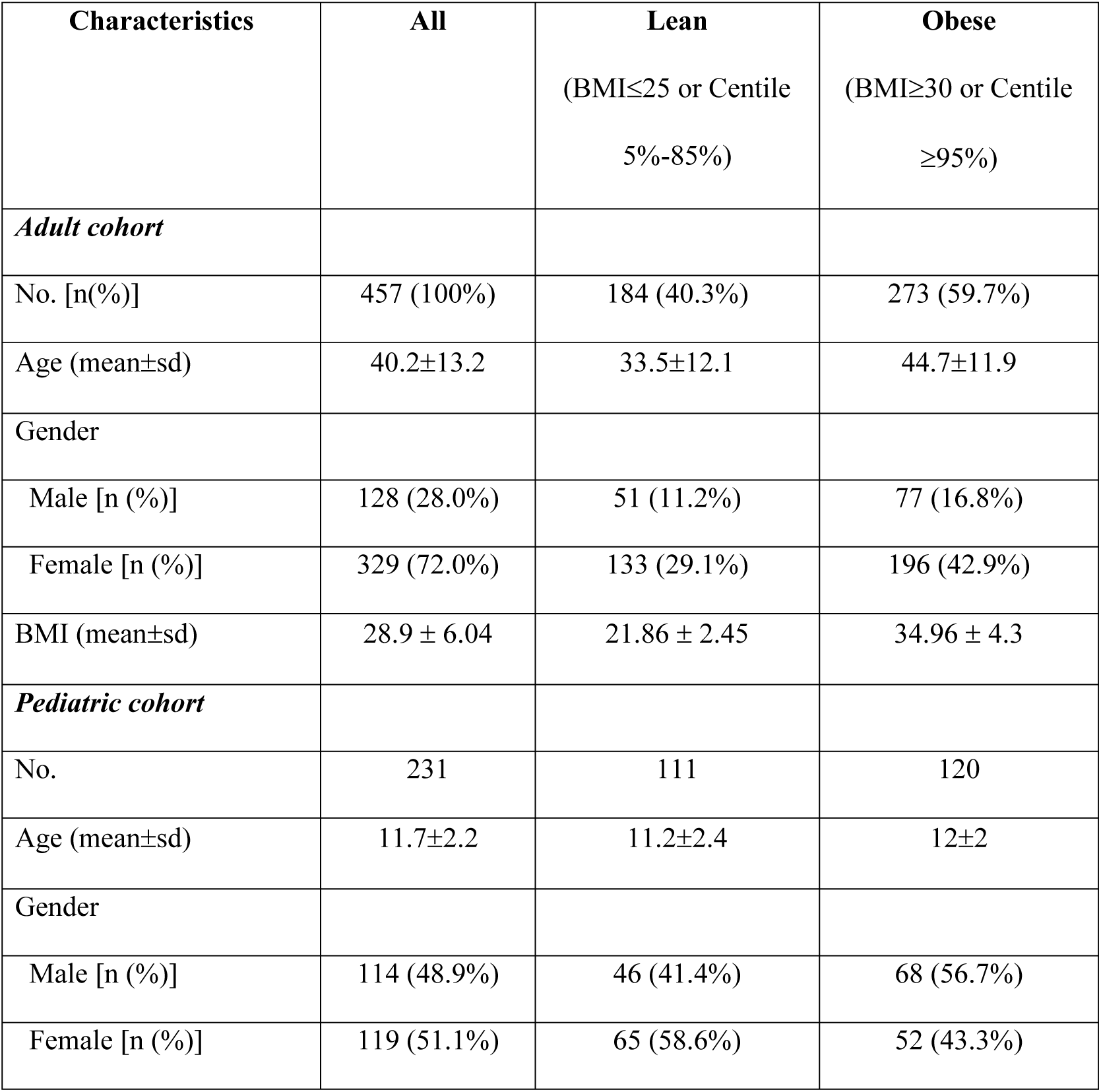
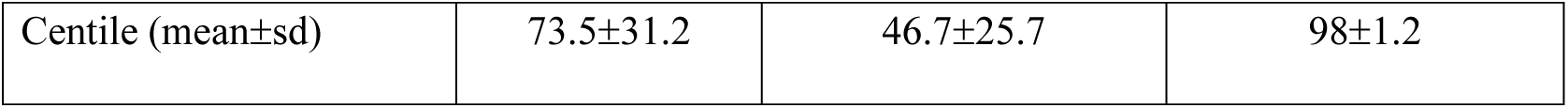
Description of the study population

| **Characteristics** | **All** | **Lean**<br>(BMI≤25 or Centile<br>5%-85%) | **Obese**<br>(BMI≥30 or Centile<br>≥95%) |
| --- | --- | --- | --- |
| ***Adult cohort*** |  |  |  |
| No. [n(%)] | 457 (100%) | 184 (40.3%) | 273 (59.7%) |
| Age (mean±sd) | 40.2±13.2 | 33.5±12.1 | 44.7±11.9 |
| Gender |  |  |  |
| Male [n (%)] | 128 (28.0%) | 51 (11.2%) | 77 (16.8%) |
| Female [n (%)] | 329 (72.0%) | 133 (29.1%) | 196 (42.9%) |
| BMI (mean±sd) | 28.9 ± 6.04 | 21.86 ± 2.45 | 34.96 ± 4.3 |
| ***Pediatric cohort*** |  |  |  |
| No. | 231 | 111 | 120 |
| Age (mean±sd) | 11.7±2.2 | 11.2±2.4 | 12±2 |
| Gender |  |  |  |
| Male [n (%)] | 114 (48.9%) | 46 (41.4%) | 68 (56.7%) |
| Female [n (%)] | 119 (51.1%) | 65 (58.6%) | 52 (43.3%) |
| Centile (mean±sd) | 73.5±31.2 | 46.7±25.7 | 98±1.2 |

### Enrichment profile of virome-wide peptide epitopes in obese versus lean population

PhIPseq data from a total of 688 specimens were analyzed. NGS read counts mapped to 115,753 peptide sequences in the VirScan library were assessed against peptide counts in the input library. On an average 586.2±190.4 and 756.1±182.9 peptides are enriched (–log*p*-value of enrichment 2.3 or higher in both technical repeats) representing 20.9±5.2 and 16.9±6.6 viral species in each of the serum specimens in the adult and pediatric population, respectively. Enrichment profile of peptides (species wise), in relation to the number of peptides in the library (per species) are not different between adult and pediatric cohorts or between obese and lean population in adult and pediatric cohorts (Figs. S1 A-C). Principal component analysis (PCA) of enriched peptides in all serum specimens categorized by adult and pediatric, and obese and lean groups shows no qualitative difference in the enrichment profile of these 4 groups, suggesting that most peptides that were enriched, were enriched in all groups (Fig. S1 D). However, average number of enriched peptides representing certain viral species such as HSV-1, HSV-2, EBV and CMV are higher (*p*<0.05) in adult obese group compared to the lean group (Figs 1 A and B). In the pediatric population, average number of enriched peptides of all of these viral species, except EBV, is higher in the obese group than the lean group. On the other hand, average number of enriched peptides of certain species of picornaviruses are lower in the obese group compared to the lean group in the adult cohort only (Figs. 1 A and B).

**Figure 1:**
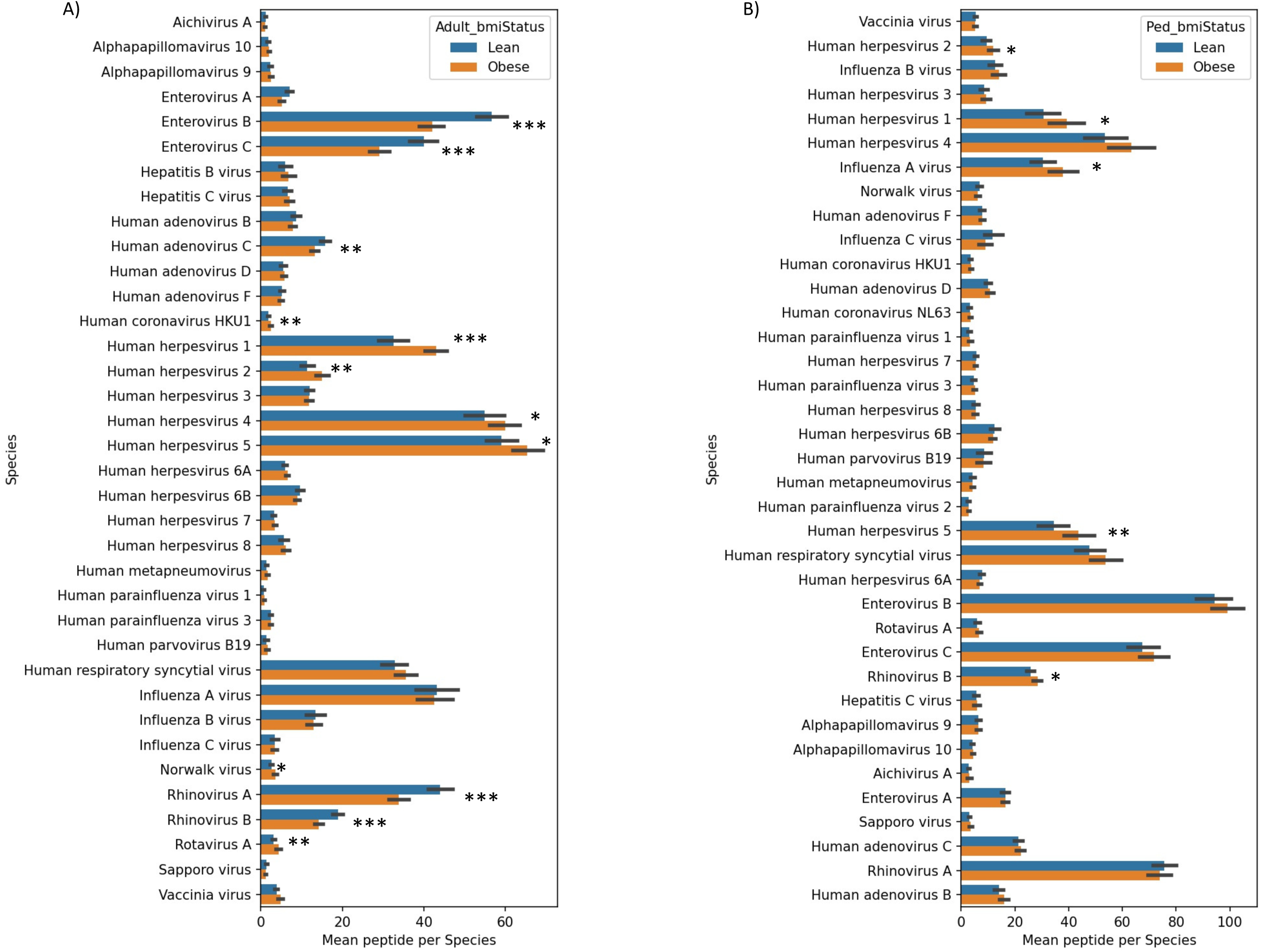
Antibody responses to viral peptides in obese versus lean populations. Peptides that were immunoprecipitated by antibodies present in the serums specimens and passed the significance and reproducibility thresholds for enrichment were counted per virus per specimen. Bar plots showing standard error mean (SEM) of the enriched peptide number for the most prevalent viruses (>10%) between the lean and obese groups of the adult (A) and pediatric (B) cohort. \**p*<0.05, \*\**p*<0.01, \*\*\**p*<0.001 by Mann Whitney’s U test.

### In-house validation of VirScan based serology data

To determine the serological status of individuals as ‘positive’ or ‘negative’ for different viruses based on VirScan data, virus-specific species score thresholds determined by a generalized linear model (GLM) was applied as described in the materials and methods. VirScan based serological data for CMV, EBV and HSV-1/2 were compared to that of conventional serology. Because our conventional HSV-1/2 IgG test does not differentiate between HSV-1 and -2, VirScan results for these viral species were combined. VirScan results either positive for HSV-1 or -2 or for both were all considered as HSV positive. Specimens from the pediatric cohort were simultaneously tested by VirScan and conventional serology. The sensitivity, specificity and accuracy of VirScan results compared to conventional methods are all 98% for CMV, 100%, 83% and 97% for EBV, and 89%, 91% and 90% for HSV, respectively (Table 2).

**Table 2:**
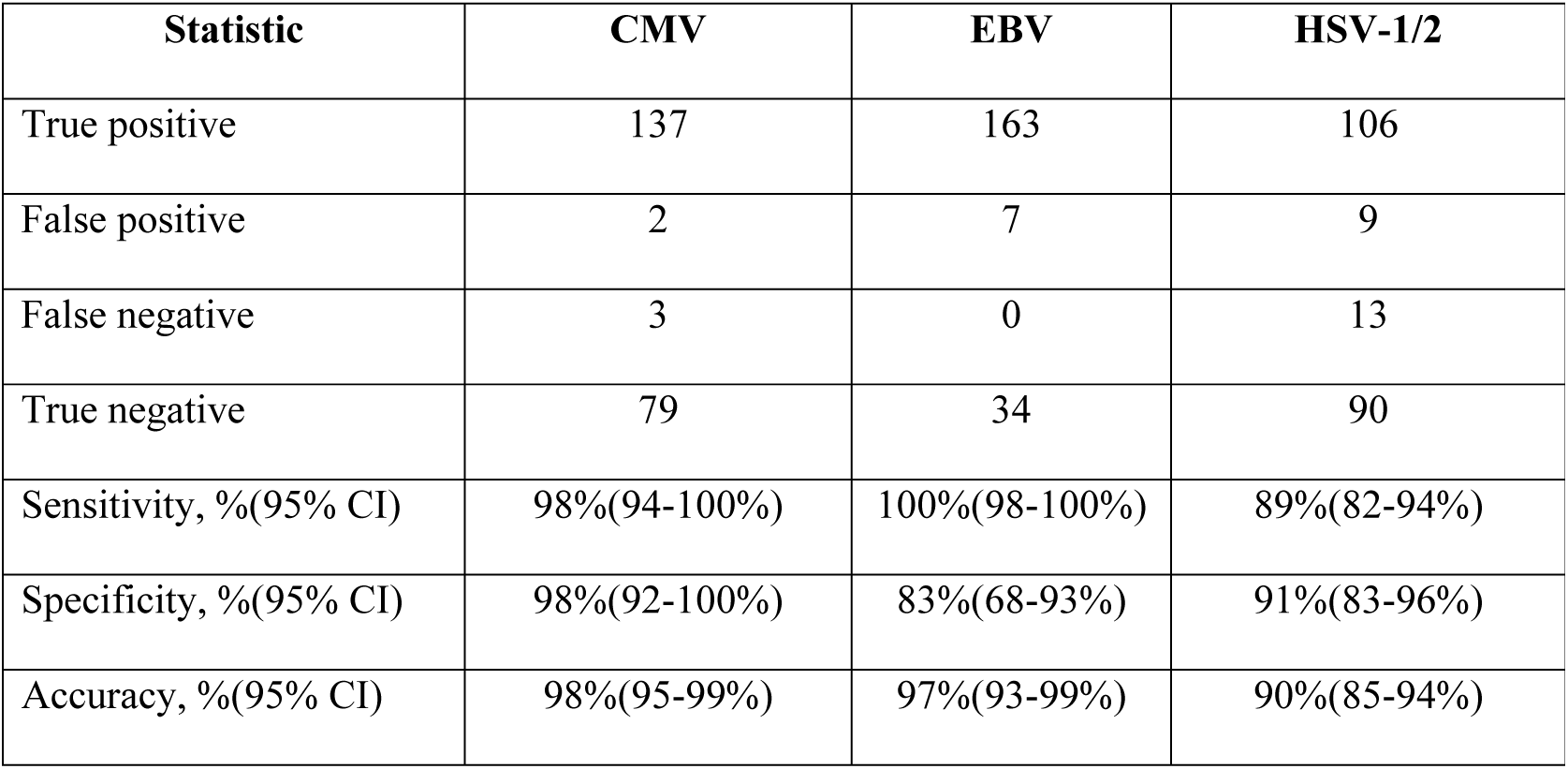
Performance characteristics of VirScan PhIP-Seq methodfor serological detection of CMV, EBV and HSV compared to standard methods

| **Statistic** | **CMV** | **EBV** | **HSV-1/2** |
| --- | --- | --- | --- |
| True positive | 137 | 163 | 106 |
| False positive | 2 | 7 | 9 |
| False negative | 3 | 0 | 13 |
| True negative | 79 | 34 | 90 |
| Sensitivity, %(95% CI) | 98%(94-100%) | 100%(98-100%) | 89%(82-94%) |
| Specificity, %(95% CI) | 98%(92-100%) | 83%(68-93%) | 91%(83-96%) |
| Accuracy, %(95% CI) | 98%(95-99%) | 97%(93-99%) | 90%(85-94%) |

### Viral seroprevalence in obese versus lean population

Virome-wide serological data based on VirScan analysis were used to compare the seroprevalence of different viral species between Qatari obese and lean populations. Among the viral species that are at least 10% prevalent in either obese or lean population in the adult cohort, seroprevalences of several members of the Herpesviridae family such as HSV-1 and - 2 and EBV are higher in the obese group and are positively associated with obesity with nominal *p*-value cut-off set at 0.05 (Figs. 2 A and B). On the other hand, seroprevalences of several members of the Picornaviridae family such as enteroviruses and rhinoviruses are lower (*p*<0.05) in the obese group and are negatively associated with obesity in this cohort (Figs. 2 A and B). However, after applying the significance threshold (Bonferroni correction, *p*<0.00115) for multiple testing, obesity among Qatari adults are significantly associated with higher odds (OR: 3.3; *p*=2.8E-08) of HSV-1 seropositivity and lower odds (OR: 0.41; *p*=0.0001) of rhinovirus A seropositivity. With the same adjusted *p*-value cut-off, no such association is observed in the pediatric population. Apart from the viral species that belong to the Herpesviridae and Picornaviridae families, the seroprevalence of rotavirus A is higher in the obese groups and is nominally associated (*p*<0.05) with obesity in both cohorts. In addition, obesity is nominally associated with higher seroprevalences of human coronavirus HKU1, human adenovirus D, influenza C virus, human parainfluenza virus 1 and human parvovirus B19 in the adult cohort and higher seroprevalences of influenza A and B viruses in the pediatric cohort. Consistent with VirScan data, seroprevalences of CMV, EBV and HSV in the pediatric cohort determined by conventional serology are not significantly different in obese versus lean groups (Fig. S2).

**Figure 2:**
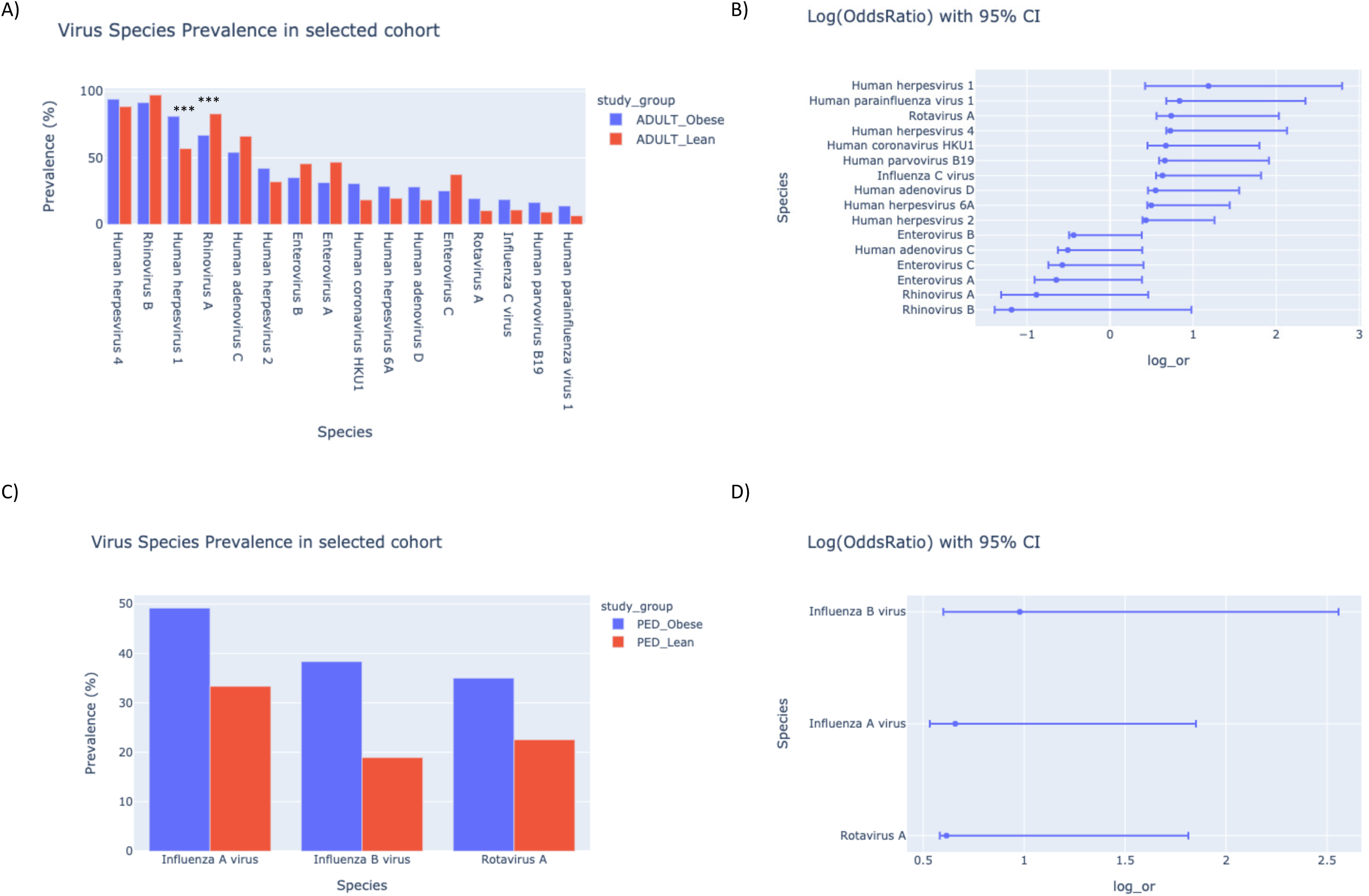
Viral seroprevalence and association with obesity. Seropositivity to a virus was determined based on adjusted species score (≥1) or total number of enriched peptides per virus normalized with the virus-specific threshold. Bar plots and forest plots show data for viruses that are at least 10% prevalent in either obese or lean or both populations and are nominally (*p*≤0.05 from Fisher’s Exact Test) positively or negatively associated with obesity. Comparison of viral seroprevalence in obese and lean population in the (A) adult cohort and (C) the pediatric cohort. ***Viruses that passed significance threshold after Bonferroni correction (*p*≤0.00115). Viral association with obesity shown by log(odds ratio) with 95% CI in the (A) adult cohort and (C) the pediatric cohort.

The seroprevalence of HSV-1 among Qatari general population is unknown but in a previous study HSV-1 seroprevalence among Qatari male blood donors was estimated to be 82.3% (19). Based on our VirScan data, the overall prevalence of HSV-1 in Qatari adult population is 74.1%, with 70.4% among the males and 76.1% among the females. The seroprevalence of HSV-1 in Qatari adult obese and lean population is 81.3% and 57.1%, respectively. Seroprevalence of HSV-1 is the pediatric population is 47%, which increased to 55% by the age of 30 years, and to 90% by the age >55 years. To assess whether our results are affected by gender and age of the study participants, we used a multivariate regression model and examined the coefficients of association with adjusted beta values. The associations of age, gender and BMI status were studied with respect to adjusted virus scores. Tests for a total of 190 associations were performed with adjusted viral scores for 38 viral species (>10% prevalence) in 457 adult and 231 pediatric samples with five features (age, male, female, lean and obese). An association was considered to be significant if absolute coefficient of association (|beta|) was ≥0.678 and *p*-value was ≤0.00013 (-log10(pval) > 3.88). A total of 25 and 4 associations were determined to be significant for the adult and pediatric cohort, respectively (Fig. 3). While in the adult cohort, HSV-1 (β = 0.739 (95%CI=0.48-0.99); -log10(p-value) = 7.82) and HHV7 (β = 0.745 (95%CI=0.562-0.93); - log10(p-value) = 14.72) are associated with obesity irrespective of age and gender, EBV is associated with female obese group (β = 1.317 (95%CI=1.05-1.5); -log10(p-value) = 21.93) and CMV is associated with male obese group (β = 0.895 (95%CI=0.66-1.12); -log10(p-value) = 13.49) only. On the other hand, rhinovirus A and B are associated with male lean group and adenovirus C is associated with female lean group only (Fig. 3A). In the pediatric cohort, no gender specific association is observed. Rhinovirus A is associated with the lean group (β = 2.07 (95%CI=1.07-3.08); -log10(p-value) = 4.27) and enterovirus A is associated with the obese group (β = 2.40 (95%CI=(1.46-3.34); -log10(p-value) = 6.29), respectively (Fig. 3B).

**Figure 3:**
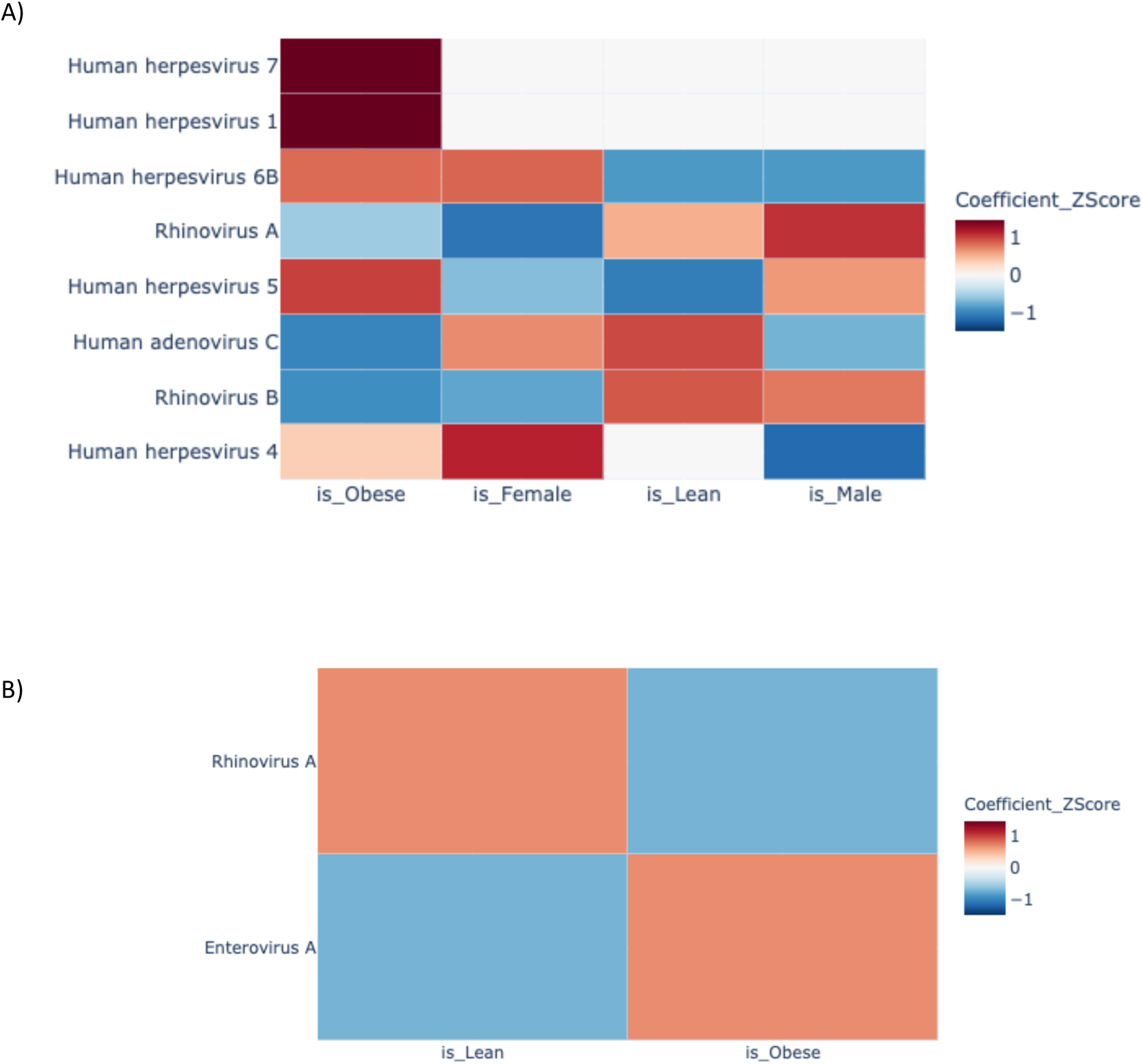
Viral association with obesity in a multivariate regression model adjusted for age and gender. The associations of age, gender and BMI status with adjusted virus scores. Heatmaps showing coefficients of association with adjusted beta values for (A) the adult cohort and (B) the pediatric cohort.

### Differentially enriched peptides in obese versus lean population

We further analyzed VirScan data to determine association of different viruses with obesity at their peptide epitope levels. Distribution of peptides that are differentially enriched in adult and pediatric obese or lean population with an odds ratio of 2 (= +/- 0.693 in log scale) with significance threshold *p*=0.005 (–log(p-value)=2.9) are shown (Figs. 4 A and 4C). The odds ratios of the same peptides grouped by viral species with 95%CI are shown in Fig. 4 B and D for the adult and pediatric population, respectively. After adjustment of *p*-values for multiple testing (Bonferroni correction, *p*<3.61E-06), a set of peptides representing members of the HSV-1 and -2 are found differentially enriched in the adult obese group, while peptides that belong to the Picornaviridae family such as enteroviruses A-C and human rhinoviruses A and B are differentially enriched in the adult lean population. With the same significance criteria, none of the peptides show significant association with obesity in the pediatric cohort. A list of HSV-1 and -2 peptides that are significantly associated with obesity and their prevalence in obese versus lean populations are shown in Table S1. A closer look at the differentially enriched (DE) peptides that are positively associated with obesity by multiple sequence alignment of each of the DE peptides in the pediatric cohort with all DE peptides in the adult cohort revealed that some peptides or potential epitopes are differentially enriched and positively associated (nominal *p-*value <0.05) with obesity in both independent cohorts. Differentially enriched antigenic peptides were primarily derived from viral structural proteins, including glycoprotein G (gG) and tegument proteins of HSV (Table S2).

**Figure 4:**
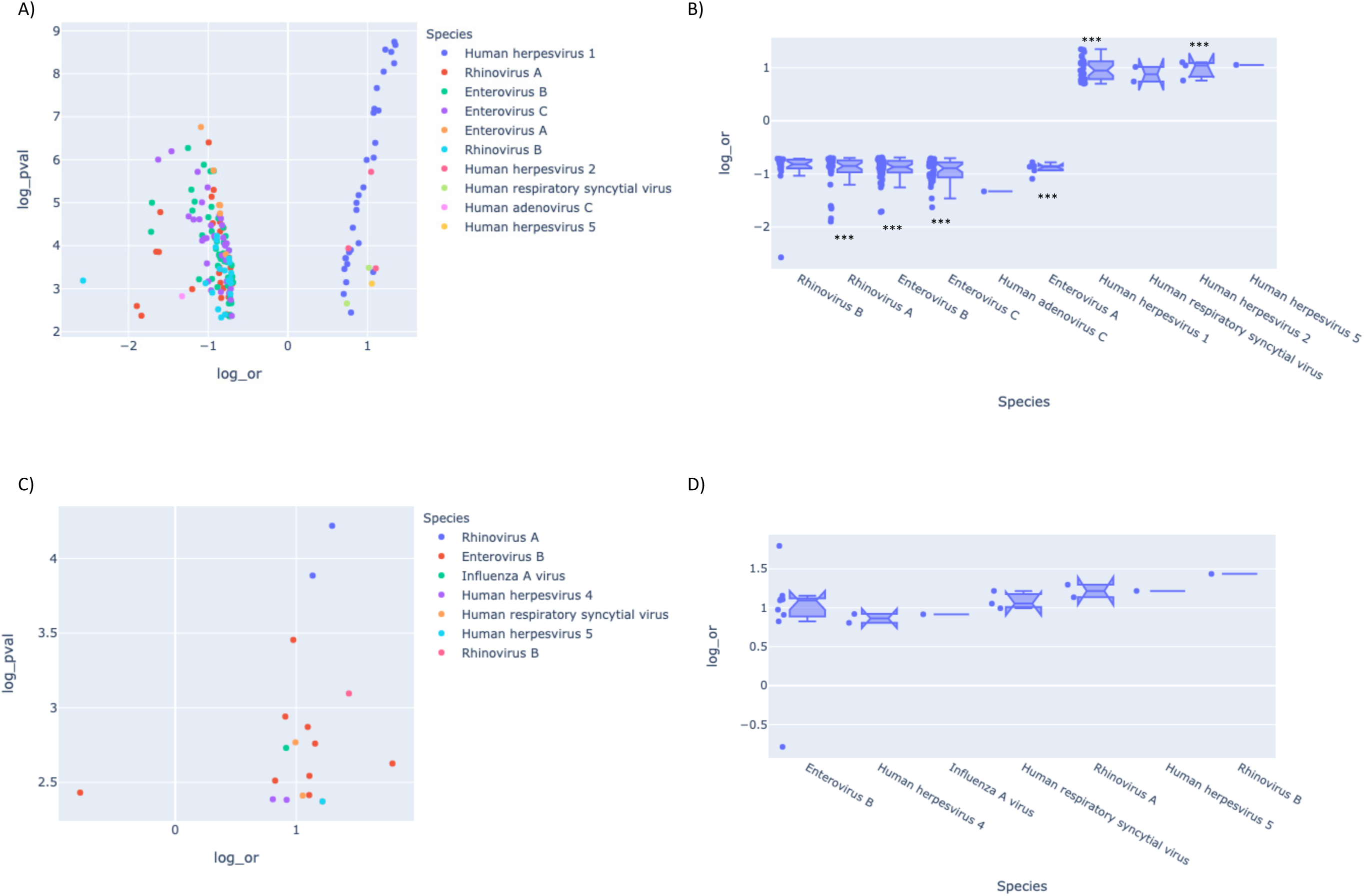
Viral peptides differentially enriched in the obese populations. Enriched peptides of the VirScan phage display library that were positively or negatively associated with obesity with a minimum odds ratio of 2 (= +/- 0.693 in log scale) with significance threshold i.e. 0.005 (= 2.9 for –Log(p-value)) are plotted. Volcano plots showing the distribution of differentially enriched peptides based on their OR and p-values in (A) the adult population and the (B) pediatric population. Violin plots showing the differentially enriched peptides grouped under viral species with their OR and 95%CI. ***Peptides that passed significance threshold after Bonferroni correction (*p*<3.61E-06).

## Discussion

Currently there is compelling epidemiological data on the association between Adenovirus-36 and obesity as well as animal data on several other viruses (7, 20-22), but a broad screen for these and other infectious causes of obesity cannot be performed without a well-established, high throughput platform for comprehensive serological profiling. The establishment of the VirScan PhIP-Seq serological profiling platform (16) has created an opportunity to investigate the association between obesity and infection at an unprecedented depth. In this study we conducted a virome-wide seroepidemiological survey with an aim to unravel associations between obesity and individual’s viral exposure history. With one of the highest rates of obesity in the world, the Qatari population represents one of the best populations for this study.

By taking advantage of the VirScan peptide library that represents the entire human virome, and its ability to characterize individuals according to their humoral immunity, we assessed interrelationships between obesity and viral infections at both species and peptide-epitope levels. For all analyses, viral species and associated peptides that have a prevalence of under 10% in either obese or lean populations were excluded. We first looked at the distribution of enriched peptides that represent the presence of antibodies specific to those peptides in the serum samples from study subjects. We also analyzed the average number of enriched peptides per virus in the obese and lean populations in both adult and pediatric cohorts. Although the distribution of enriched peptides is not dissimilar between different group of subjects (Fig. S1), and the the average number of enriched peptides of most viral species are not significantly different in the obese versus lean population, average number of enriched peptides of herpes viruses are significantly higher (*p*≤1.0E-05 for HSV-1) in the obese groups compared to the lean groups. On the other hand, the average number of enriched peptides of several viral species of picornaviruses are higher in the lean population compared to the obese population in the adult cohort only. While these results suggest the presence or absence of additional features in the antibody profiles of obese individuals directed towards these viruses, in order to study the association of obesity with viral exposure history, we determined the seroprevalence of these viruses in Qatari obese and lean populations.

Unlike previous VirScan-based studies that applied empirically determined virus-specific species scores to determine seropositivity (16, 18), we established a generalized linear model using control serum, tittered for antibodies against different viruses, and taking into the accounts of number of peptides available in the library for each viral species. We then tested the specimens from the pediatric cohort for CMV, EBV and HSV with standard serological assays that are used for patient testing. Serological data on HSV, EBV and CMV were then used to validate the VirScan-based serological data demonstrating 90% to 98% accuracy compared to the standard methods. Based on VirScan based serology, after adjustment of *p*-values for multiple testing, HSV-1 seropositivity is significantly associated with obesity in the adult population only (Figs. 2 A and B). Consistently, HSV-1 is associated with obesity in the adult population independent of age and gender by multivariate regression analysis (Fig. 3A). Other herpesviruses such as HSV-2, CMV and EBV are also nominally associated with obesity, and by multivariate analysis, they are associated with obesity in a gender specific manner.

To obtain further insights on the relationship of herpes viruses with obesity, we analyzed the association of virus specific peptides with obesity. Consistent with our findings at the species level, it is mostly the HSV-1 associated peptides that show the strongest associations with obesity (Fig. 4 A and B). Peptides from other herpes viruses are also nominally associated with obesity in the Qatari adult population. Seroprevalence of herpesviruses are not significantly different between obese and lean population in the pediatric cohort but differentially enriched, HSV-1/2 and EBV peptides are nominally associated with obesity (data not shown). We also found some HSV-1/2 peptides that are independently associated with obesity in both populations (Table S2). These results suggest that acquiring HSV-1 infection as people gets older may increase the odds of becoming obese. An early sign of this is observed by the nominal association of HSV peptides with obesity in the pediatric population. Positive association of these HSV-1/2 peptides with obesity observed in our population also correlates with earlier evidence of association between HSV and excessive adiposity in different age and gender groups. Our data on differentially enriched peptides in the adult population is consistent with the cross-sectional data from National Health and Nutrition Examination Survey (NHANES) in USA, during the period of 1999–2012, showing significantly higher prevalence of HSV-1 in both obese men and women (23). In another NHANES survey in 1999-2004, CMV was found to be significantly associated with high BMI in the female population only (ages 20-49 years). However, in our adult (>18 years) population, seropositivity to CMV was associated with obesity in the male population only.

In our study, obesity was not associated with human adenovirus or any of its serotypes (Adv-5, Adv-9, Adv-31, Adv-36 and Adv-37) that were previously reported to be associated with obesity, epidemiologically, or linked to enhanced lipogenesis *in vitro*, suggesting that adenoviruses may have a limited role in the incidence of high rates of obesity among Qataris. In a recent study, Lessan *et al*. demonstrated that the seroprevalence of Adv-36 in United Arab Emirates (UAE) is much higher in their population although there is no significant difference in prevalence of this virus in their obese and lean populations (24). Our adenovirus data provides additional support to this finding in a highly similar population.

An incidental but interesting finding in our study is the negative association of the family of picornaviruses with obesity. This phenomenon was only observed in the adult population and may potentially be related to the waning immunity to these viruses with age. Obesity associated impaired immunity to various infections such as influenza, pneumonia, *Helicobacter pyrolii* and nosocomial infections or poor response to vaccines such as hepatitis B, tetanus and influenza vaccines have been described in the literature (21). However, there are no reports on impaired immunity or higher susceptibility to infections with picornaviruses such as enteroviruses and rhinoviruses in relation to obesity. Interestingly, no such association is seen in the pediatric population. This may be related to higher rates of infection with these viruses in children, and their immune systems being continuously challenged by these viruses. These findings are novel and warrant further investigations in a separate study.

Using a comprehensive virome-wide analysis, we identified viral species previously associated with obesity, but were unable to find novel viral associations with obesity. However, the association of different herpesviruses or their specific epitopes with obesity confirms the association of these viruses described for other populations. Furthermore, despite very high rates of obesity, no viral association studies have ever been described for the Qatari population. One limitation of our study is that VirScan results are based on antibody binding of linear epitopes only. Therefore, potential antibody interactions that relies on tertiary structure of epitopes may have been missed. Also, our clinical validation of VirScan serology data was limited to CMV, EBV and HSV only. Our study failed to provide any mechanistic insights on the role of HSV-1 infection in adipogenesis. But, for the first time, we have described high resolution, peptide-epitope level data for HSV-1 in association with obesity. Our analysis of differentially enriched peptides from a large collection of peptides covering the entire human virome reveals a number of viral epitopes that could be utilized for further mechanistic studies. We have listed candidate HSV-1 and -2 peptides that were strongly correlated to obesity in the adult population and are highly prevalent in both adult and pediatric obese populations (Table S 1 and 2). Interestingly, the majority of these peptides belongs to the HSV glycoprotein family. Both HSV and CMV are known as lipogenic viruses and are known to affect cellular metabolism in different ways or are known to cause expansion of adipose tissues by their effects on inflammatory pathways (7, 10). Further studies on the role of HSV-1 candidate peptides identified in this study in cell culture or animal models may reveal novel pathways for virus induced adipogenesis.

In conclusion, we have conducted a virome-wide seroprevalence study to detect associations between past viral exposures and obesity in a population that is highly endemic for obesity. Our analysis revealed a strong positive association of obesity with HSV-1 and nominal associations with other herpesviruses. Our finding may have implications for understanding the underlying causes of higher prevalence of obesity not only among the Qataris but also among the citizens of other countries in the Arabian Peninsula.

## Materials and Methods

### Study design and subjects

For the adult cohort, 800 randomly selected serum specimens were obtained from Qatar Biobank (QBB). This cohort represents Qatari nationals and long-term residents (lived in Qatar for >15 years) aged ≥18 years. Data on age, gender, ethnicity and BMI were collected. Specimens from non-Qatari participants and specimens for which no BMI data were available were excluded from analysis. Also, specimens from subjects that fall into the overweight category (BMI >25 to <30) were excluded from the analysis. For the pediatric cohort, 231 serum samples were collected from Qatari obese and lean children admitted to Sidra Medicine, which is a 400-bed tertiary care children’s and women’s hospital in Qatar, during the period October 2018 to November 2019. Residual specimens from comprehensive metabolic panel (CMP) and basic metabolic panel (BMP) tests were collected. Specimens from subjects that satisfies inclusion criteria were identified through laboratory information management systems (LIMS) query on a weekly basis using Discern Analytics 2.0 (Cerner). The inclusion criteria were: i) age 7-15 years ii) Qatari nationality and, iii) BMI centile 5% to 85% as lean individuals or ≥95% as obese. Underweight (BMI centile <5%) or overweight (BMI centile 85% to <95%) children were excluded. Also, children with chronic diseases such as Cancer, Type 1 Diabetes, Immunosuppression, developmental delay and children with history or recent infections were excluded. Data on age, gender, ethnicity and BMI were collected. Specimens were aliquoted and stored at -80°C until further tested. Ethics approval for the study was obtained from the Institutional Review Boards of Sidra Medicine and Qatar Biobank.

### VirScan phage immunoprecipitation sequencing (PhIP-Seq)

VirScan PhIP-Seq analysis of serum specimens from the adult and pediatric cohort was carried out by the methods described previously except that an expanded bacteriophage library (containing 2×10^10^ plaque-forming units) displaying 115,753 peptides was used, and sequencing was performed using an Illumina NextSeq500 platform and NextSeq 500/550 High Output Kit v2.5 (75 Cycles) kits (Illumina). Each specimen was tested in two technical replicates. Each sequencing batch consisted of specimens representing obese and lean subjects, mock-IP controls, positive control and input library in a random manner with an aim to minimize batch effects on the output data. PhIP-Seq analysis of a total of 688 specimens was completed in 12 NextSeq500 sequencing runs.

### Serological assay

Serum specimens from the pediatric cohort were assessed for IgG antibody titers against CMV (n=221), EBV (n=221) and HSV-1&2 (n=219) using LIAISON® CMV IgG II, EBNA IgG and HSV-1/2 IgG assays on a LIAISON® XL automated chemiluminescence analyzer (DiaSorin) according to manufacturer’s instruction and standard operating procedures (SOP) at the Serology Laboratory of Sidra Medicine. The results were interpreted as positive, negative and equivocal according to manufacturer’s criteria. Equivocal results were excluded from analysis.

### Statistical analysis

Descriptive statistics were presented as means ± standard deviation (SD) for continuous variables or as numbers and percentages for nominal/categorical variables. VirScan data were filtered for enriched peptides as described previously (16). Briefly, we first imputed *p*-values (-log10 transformed) by fitting a zero-inflated generalized Poisson model to the distribution of output counts followed by regressing the parameters for each peptide sequence based on the input read counts. Peptides with a reproducibility threshold (-log10(P-value) ≥ 2.3) in both technical repeats were further filtered for sporadic hits by removing peptides which were also significantly enriched in at-least two mock-IP controls (beads only). We also considered a peptide hit to be significant if the same peptide enriched in more than one sample. Average number of enriched peptides for different viral species were compared using *p*-values from Mann-Whitney U test for each species.

We counted the number of non-homologous (i.e. peptides that do not share more than seven linear sequence identity), enriched peptides for each virus to evaluate virus specific scores, which has linear relation with virus peptidome size, i.e. number of available peptides in the input library. To mitigate this effect, we evaluated an adjusted virus threshold score with a generalized linear model (GLM), in which we regressed the virus library size and number of enriched peptides to model virus specific cutoff from known in-house cases as described earlier (25). In brief, specimens with known IgG titers for CMV, EBV, HSV, Varicella Zoster, Mumps, Measles, Rubella, Hepatitis A, Hepatitis B, and Parvovirus B19 viruses were used. We extrapolated the adjusted virus-specific scores using this model, which has been further used for seroprevalence calculations. We compared the seroprevalence at virus species level using adjusted virus score. Different group-wise comparisons were performed to compute Odds Ratios (OR) and *p*-values from Fisher’s Exact test with Bonferroni correction for multiple testing. We considered a virus to be differentially enriched in one group if the prevalence is ≥10% in at least one group, |log(OR)| ≥ log10(2) and *p*-value ≤ 0.05/n; where n is the number of tests (i.e. virus species).

Multivariate logistic regression analysis was performed to test for associations of age, gender and BMI status with adjusted virus scores (after virus specific thresholds were applied) instead of categorical results. An adjusted *p*-value <= 0.005/n and |beta| ≥0.678 were applied to filter data for most significant results. Finally, differentially enriched peptides in obese versus lean populations were determined based on prevalence, OR and *p*-values from Fisher’s Exact test, corrected for multiple testing (Bonferroni correction). Similar thresholds were applied (prevalence ≥10% in at least one group; |log(OR)| ≥ log10(2) and *p*-value ≤ 0.05/n). All statistical analyses were performed using python (v.3.7) with statistical modules statsmodel (v.0.11.1) and sklearn (v.0.22).

## Acknowledgments

We would like to thank the Qatar Biobank (QBB) management and staff, for their time and effort allowing us to access and analyze samples and data from the Qatar Biobank, and Dr. Stephan Elledge (Brigham and Women’s Hospital and Harvard Medical School, Boston, MA) for kindly providing the VirScan phage library used in this study.

